# Development of a Microdroplet-Based Functional Genomic Screening pipeline by combination of DNA Nanoflowers and PURExpress Cell-Free Expression

**DOI:** 10.64898/2026.01.27.701873

**Authors:** Joan Castells-Ballester, Abigail Taron, Madison Smith, Rebecca Gawron, Julie Beaulieu, Olta Papa, Jackson Buss, Jennifer Ong, Minyong Chen

**Affiliations:** New England Biolabs

**Keywords:** Cell-free expression, Functional Metagenomics, Microdroplets, Microfluidics, Nanoflowers

## Abstract

We present a microfluidic workflow that couples reconstituted in vitro transcription–translation (IVTT) with ultra–high-throughput droplet screening to directly link genotype and phenotype within complex, heterogeneous DNA pools. The approach employs DNA nanoflowers as clonal, high-copy templates, enabling robust protein expression from single DNA molecules encapsulated in picoliter droplets. When integrated with fluorescence-assisted microdroplet sorting (FADS) and a DNA recovery pipeline that reconstituted selected libraries for subsequent iterative rounds, the platform achieves approximately 400–500-fold enrichment per selection cycle and supports functional discovery and directed evolution entirely independent of host cell expression.

As a proof of principle, we demonstrate recovery of the recombinase RecA from an *E. coli* genomic library screened for single-stranded DNA binders, highlighting the platform’s capability to identify DNA-interacting and DNA-modifying enzymes. By eliminating host-derived background activity and toxicity constraints that often complicate lysate- or cell-based metagenomic screens, this method expands access to enzyme classes that have historically been difficult to assay.

## Introduction

Compartmentalizing biological reactions is a natural core strategy that living systems use to stay organized and efficient. By separating different processes into defined spaces, it is possible to fine-tune how and when reactions occur. In the field of protein engineering and discovery, enzymatic activities derived from individual genetic variants are confined and evaluated in separate compartments, allowing researchers to directly observe the functional consequences of each genotype through its biochemical output. This phenomenon is referred to as genotype-phenotype linkage and occurs naturally when analyzing biological systems, such as individual cells, or when cellular material is analyzed as a readout well in a microtiter plate.

Over the past decades, droplet microfluidics has emerged as a powerful technology for establishing genotype– phenotype linkage while drastically reducing reaction volumes and screening costs (reviewed in Gantz et al., 2023). When combined with a measurable output of enzyme activity, such as fluorescence or colorimetric substrate conversion, this platform enables ultra-high-throughput screening with unprecedented scalability. Droplet microfluidics has been used to increase enzyme activity and/or modify enzyme specificity in protein evolution campaigns (Agresti et al., 2010; Holstein et al., 2021; Ma et al., 2018; Penner et al., 2025) and discover new enzymes in functional metagenomics workflows (Alma’abadi et al., 2022; Colin et al., 2015; Gantz et al., 2023; Neun et al., 2022a; Olagnon et al., 2024; Tauzin et al., 2020).

Traditionally, functional metagenomic screening is microtiter plate-based, and it involves cloning environmental DNA fragments into fosmid vectors, which harbor several kilobases of eDNA sequence, and expressed in *E. coli* cells using endogenous transcription and translation machinery. Cell lysates from the individual colonies grown in microtiter plates are then assayed to identify clones with the desired activity (Chuzel et al., 2018; Leis et al., 2013). Similarly, the vast majority of functional metagenomic approaches using microfluidics developed in recent years exploit microbial hosts (mostly bacterial) to express the target enzymatic activity and are encapsulated into droplets at the single-cell level in the presence of a reporter fluorescent substrate. Release of the target catalyst is then achieved by In-droplet cell lysis and; and positive droplets are usually sorted by fluorescence-assisted microdroplet sorting (FADS) or comparable methods for DNA recovery and sequencing (Asama et al., 2024; Colin et al., 2015; Gielen et al., 2018; Nakamura et al., 2025; Neun et al., 2022b). These approaches work well for many enzymes but rely on assays performed in cell lysates, which can cause two main problems. First, endogenous enzymes such as nucleases and proteases from the host can react with the substrate, producing high background and false positives. Second, lysates contain components that could potentially interfere with the target functional assay, create noise, or change enzyme behavior, lowering assay sensitivity. Enzyme families that are particularly challenging to screen in lysates include broad-range proteases, specific nucleases, ligases, recombinases, deaminases, and other nucleic-acid–interacting enzymes. Overexpression of many of these targets also causes host toxicity, which biases library representation and reduces the chance of recovering true positives.

An attractive approach to overcome these limitations and enhance the effectiveness of functional metagenomics is the use of fully reconstituted cell-free expression systems either ‘home-made’ or commercially available such as PURExpress (New England Biolabs, NEB) or PureFlex (GeneFrontier). These systems contain purified components required for *in vitro* transcription and translation (IVTT) (Shimizu et al., 2001), resulting in very low levels of nucleases and proteases. This minimal background enables highly sensitive functional assays. These systems rely on the encapsulation of single copy genes in microdroplets followed by expression and functional assay and have shown success in several studies (Fallah-Araghi et al., 2012; Holstein et al., 2021; Sierra et al., 2022).

However, two main limitations hamper the application of cell-free in metagenomics. First is that expressing detectable amounts of protein from a single DNA copy in a picoliter volume is often insufficient for activity assays (Holstein et al., 2021; Sierra et al., 2022); therefore, DNA must usually be amplified or present at high copy number in each droplet to produce enough protein for reliable functional screening.

Isothermal DNA amplification with Rolling Circle Amplification (RCA) methods present an attractive, amplicon length-independent option to be combined with IVTT for cell-free protein expression (Kumar & Chernaya, 2009). But typically, combining DNA amplification and IVTT using reconstituted transcription-translation systems in one-pot reaction is not possible due to incompatibility between the different reaction components (Holstein et al., 2021; Seo & Ichihashi, 2023). Although RCA-IVTT alternative methods have been reported (Abil et al., 2023), these are two-step approaches in which DNA amplification is sequentially followed by IVTT and require specialized and complex microfluidics systems to merge or inject preexisting microdroplets with new reagents in segmented workflows (Holstein et al., 2021). Notably, under optimized conditions, RCA products can self-assemble into flower-like nanoparticles (“DNA nanoflowers”) through complexation of amplified DNA with magnesium pyrophosphate, a precipitate formed as a byproduct of RCA. These DNA nanoflowers enable convenient isolation and quantification of RCA products and have already been shown to be compatible with microfluidic workflows (Galinis et al., 2016; Zubaite et al., 2017). Alternatively, it is possible to generate clonal DNA on beads by solid-phase PCR in a first emulsion to be re-encapsulated into a second emulsion containing IVTT reagents (Fallah-Araghi et al., 2012; Kojima et al., 2005). The viability of this approach is limited to amplicon lengths of some kilobases and reduces the number of viable IVTT droplets to ∼10-15% due to the Poisson distribution.

The second limitation to integrating IVTT into microdroplets for functional metagenomics is that metagenomic libraries contain unknown DNA fragments of varying lengths and heterogeneous base composition, which hinders DNA recovery and prevents reliable expression of functional open reading frames that typically require defined, controllable promoters.

In this work we describe a novel workflow that integrates IVTT protein expression in microdroplets with DNA expression and recovery from complex, heterogeneous DNA pools. We combine DNA nanoflowers as clonal templates for IVTT expression with an efficient, ultra-high-throughput FADS selection module and a recovery pipeline that reconstitutes selected libraries for iterative rounds of selection. As a proof of principle, we retrieved the recombinase RecA from an *E. coli* genomic screen for single-stranded DNA binders. We envision this workflow overcoming the limitations of plate-based and *in vivo* functional metagenomics and substantially expanding the catalog of enzymatic activities accessible to enzyme discovery.

## Materials and Methods

### Materials

#### Plasmids and bacterial strains

For all cloning procedures, including metagenomic and genomic library construction, plasmid reconstitution by HRR (results Section 2), either NEB^®^10-beta High Efficiency Competent *E. coli* (C3019) or NEB^®^ 10-beta Electrocompetent *E. coli* were used (C3020). For protein expression, NEB^®^ T7 Express High Efficiency Competent *E. coli* (C2566) was used.

pET-T7-sfGFP, pET-T7-mCherry, pUCT7-sfGFP, JJB668, JJB270, pMC174, pMiniT2.0 (included in NEB’s cloning kit # E1202) were made using NEBuilder HiFi DNA Assembly Cloning Kit (NEB), and gene/DNA oligonucleotide synthesis was provided by IDT (Coralville, IA). Plasmid sequences are available upon request.

#### Analysis of Nucleic acids

DNA derived from gDNA extraction, DNA fragmentation, plasmid preparation, endonuclease digests, PCR or RCA was analyzed by standard gel electrophoresis using 1% agarose.

When indicated, DNA was analyzed using 4200 Tape Station System (Agilent, CA, USA) and the Genomic DNA Screen Tape Analysis kit following manufacturer’s instructions.

#### DNF generation and purification

To generate DNA nanoflowers (DNFs, described in figure 2A), the plasmid template was diluted to achieve the desired copy number per droplet (λ1), based on an estimated maximum microdroplet diameter of 50 μm (Figure S4) and a total emulsion volume of 200 µl (∼4 x 10^6^ microdroplets). Under these conditions, a λ1 value of one is met with ∼4 x 10^6^ plasmid copies, which for an average library size of 3 kb corresponds to 30 pg of DNA.

**Figure 1.**
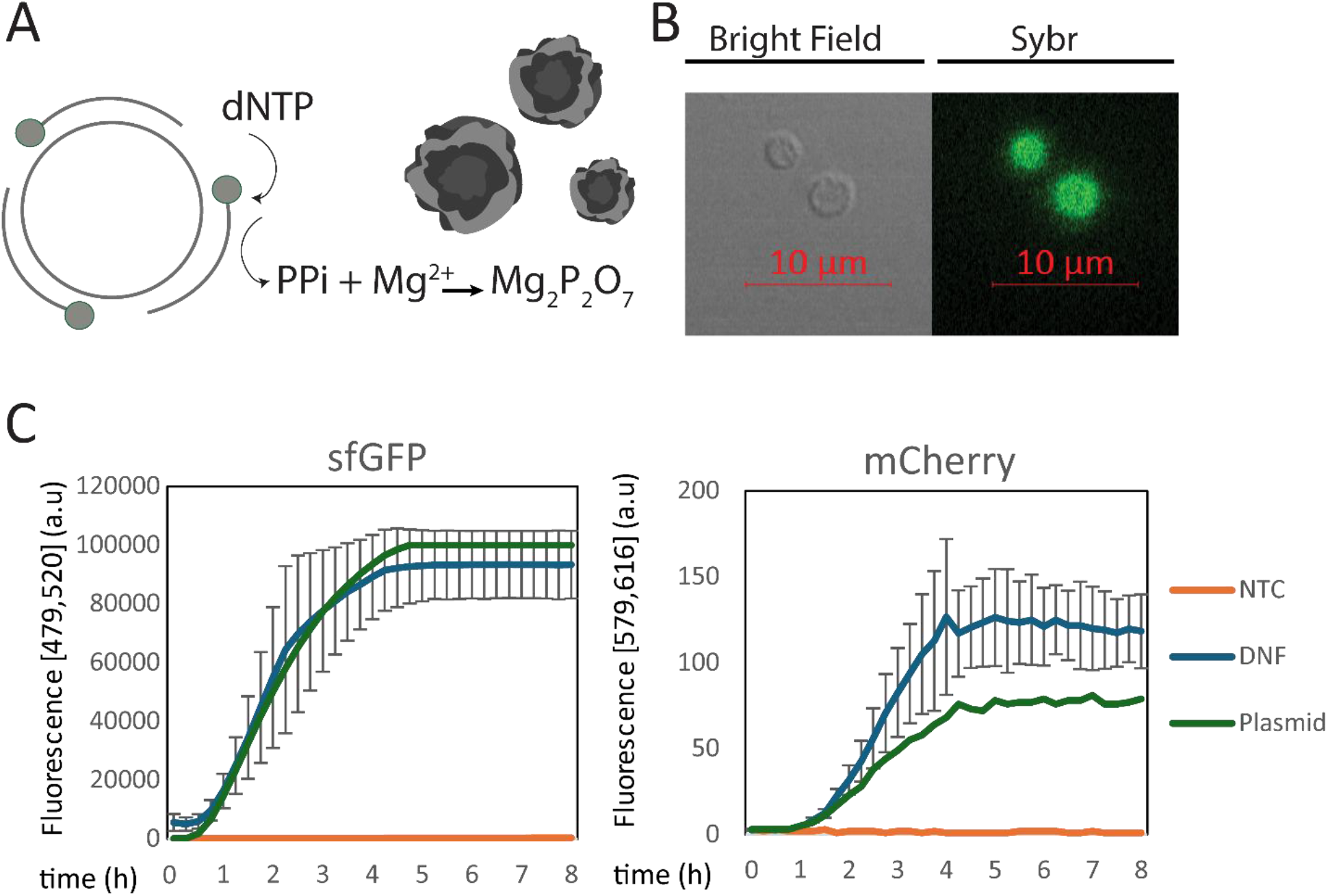
DNF is fully compatible with *in vitro* transcription and translation. (**A**) Illustration of DNA nanoflowers (DNFs) generated by self-assembly of DNA produced via rolling circle amplification (RCA). (**B**) DNFs stained with SYBR Gold. (**C**) DNFs produced from plasmids encoding sfGFP (pET-T7sfGFP) or mCherry (pET-T7mCherry) were used as templates for in vitro transcription–translation (IVTT) using PURExpress. For comparable protein expression, 5.60 × 10^7^ plasmid copies or 6.50 × 10^3^ purified DNFs were used as templates. Fluorescence at the sfGFP and mCherry wavelengths was recorded in a plate reader over an 8-hour IVTT time course.

**Figure 2.**
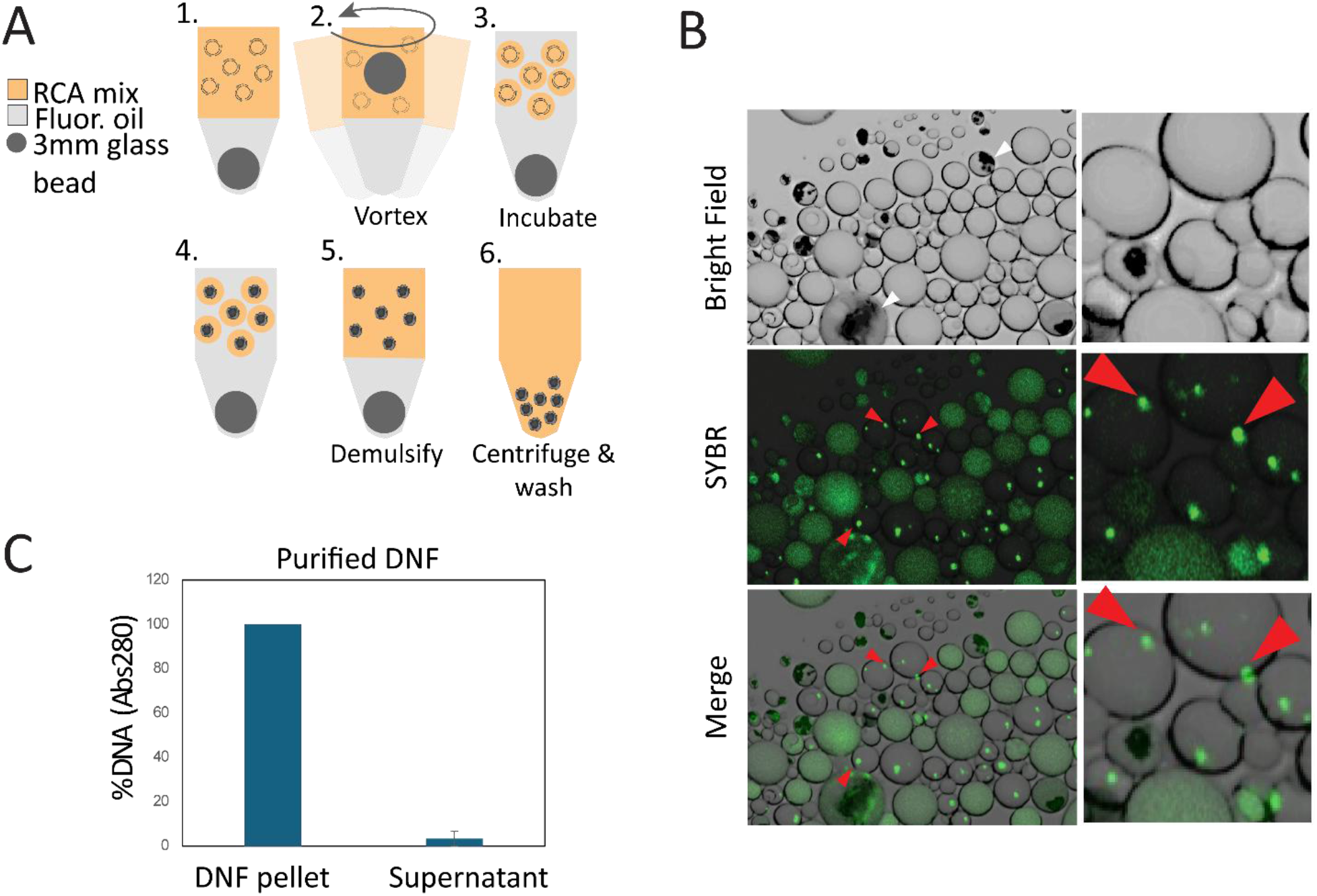
(**A**) Schematic microfluidics-free generation of clonal DNF in water-in-oil microdroplets (see methods). (**B**) Polydisperse RCA emulsion after incubation shows the presence of 1-2 DNF per microdroplet (indicated with red arrows). DNA resulting from non-specific RCA or not effectively incorporated into DNFs is shown with white arrows. (**C**) Abs_280_ measurement of the resulting purified DNF suspension before and after centrifugation shows that the vast majority of DNA remains insoluble and as part of DNFs.

The diluted template was then mixed with all components of the phi29-XT RCA kit from NEB (#E1603) in a 200 ul reaction, in a 1.5 ml tube. The RCA components were mixed following the manufacturer’s instructions with the following modifications: the standard reaction buffer was replaced with the buffer described by Galinis et al. (2016), consisting of 33 mM Tris-acetate (pH 8), 10 mM Mg-acetate, 66 mM K-acetate, 0.1% (v/v) Tween-20, and 1 mM DTT. Additionally, 0.4% (w/v) Pluronic F-127 was added to the reaction mixture, and the reaction mix was kept on ice to prevent premature amplification.

As the oil phase, HFE7500 (3M™ Novec™ 7500 Engineered Fluid) was supplemented with 3% 008-fluorosurfactant from RAN biotechnologies (Beverly, MA, USA) and a total of 200 µl was added to the RCA reaction mix. At this point, a 3 mm glass bead (Millipore Sigma, MA, USA) was added to the tube. The mix was then vortexed at 3000 rpm for 30 seconds using a Vortex-Genie 2 (Scientific industries Inc. NY, USA). The resulting emulsion was chilled on ice for 1-2 minutes and incubated at 37°C for 10 hours for DNA amplification.

After incubation, the emulsion was destabilized by adding 300 ul of PFO (1H,1H,2H,2H-perfluoro-1-octanol, Millipore Sigma, MA, USA). After briefly vortexing and a quick spin, two clear phases including a translucent precipitate were clearly visible. The upper aqueous phase was recovered and transferred to a new tube and water was added to a final volume of 1 ml. The tube was centrifuged at 8000x *g* for 8 minutes, generating a white-translucent pellet containing DNFs. After removing the supernatant, the pellet was resuspended in 1 ml of water by pipette action to wash the DNF and remove remaining RCA components. Washing was repeated 3 times. After that, DNFs were resuspended in 50 ul of water using a 21G-needle (Beton Dickinson, NJ, USA) to ensure proper resuspension.

To estimate the DNF numbers, DNF were diluted 20 times and stained using 10X SYBR gold (#S11494, ThermoFisher Scientific, MA, USA) and counted (Figure S4) using a hemocytometer under a Zeiss ApoTome Axiovert 200M microscope equipped with AxioCam MR camera.

#### Generation of monodisperse IVTT microdroplets

Monodisperse water-in-oil microdroplets were generated using the µEncapsulator System (Dolomite Microfluidics, UK), using a 2 Reagent 50 um T-junction fluorophilic microfluidic chips (#3200445). As a carrier-oil phase, HFE7500 (3M™ Novec™ 7500 Engineered Fluid) supplemented with 3% 008-fluorosurfactant from RAN biotechnologies, was used.

As IVTT reconstituted system the PURExpress^®^ *In Vitro* Protein Synthesis Kit from NEB (MA, USA) was used following the manufacturer’s instructions and functioned as the aqueous phase during microdroplet generation. In total, 200 µl including the DNFs suspension diluted to reach the desired DNF/droplet (λ2, see figure 3A and 5A) was loaded as 100 ul in each chamber of the sample reservoir.

**Figure 3.**
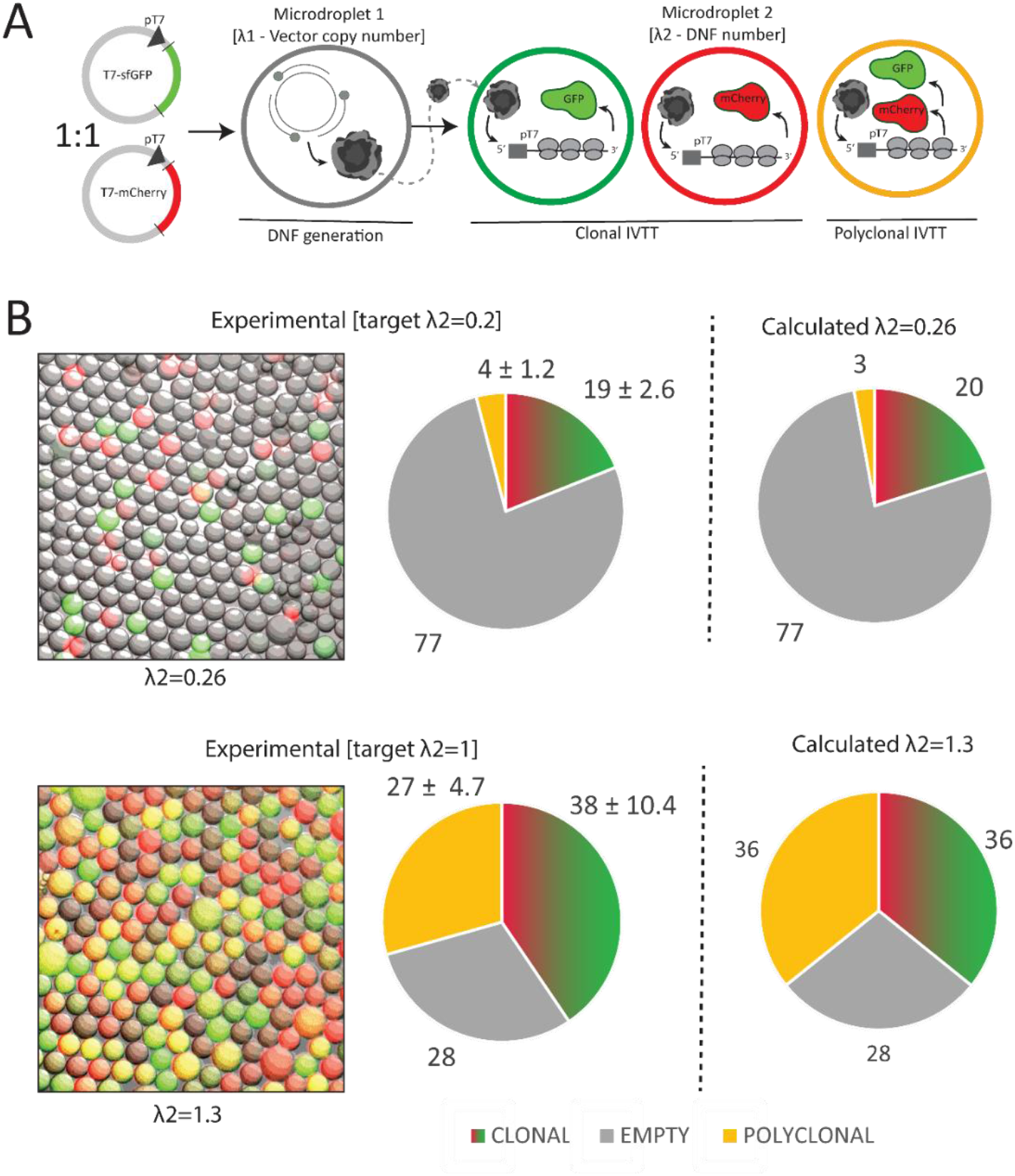
(**A**) Two-step workflow design to assess DNF clonality. An initial emulsion produces DNF from single copy vectors expressing either GFP (pJJB270) or mCherry (pJJB668) (λ1=0.5). After purification, DNF are re-encapsulated into IVTT droplets targeting λ2=0.2 or λ2=1.3, producing Green (clonal), Red (clonal), Yellow (polyclonal) or empty (no expression) (**B, left**) Confocal microscopy pictures of IVTT reactions encapsulated in 50 µm microdroplets. (**B, right**) Comparison of experimental and calculated distribution values adjusted to each corresponding λ2 according to the Poisson distribution.

Monodisperse 30-50 um microdroplets were generated at controlled flows: 10 ul/min for the carrier-oil phase and 4–5 ul/min for each aqueous phase loaded in each 100 ul sample chamber. During encapsulation, temperature was kept at 4°C using either ice-cold metal blocks or the TCU-100 temperature control unit (Dolomite microfluidics, UK). At these conditions, the 200 ul emulsion was completed in approximately 20–30 minutes. After that, the emulsion was incubated for a minimum of 10h to trigger protein synthesis.

To perform the genomic screen for single-stranded DNA (ssDNA) binders or single-stranded DNA nucleases (see Results, Section 4), we used the PURExpress® ΔRF123 Kit (NEB, #E6850S, MA, USA). All reactions were carried out following the manufacturer’s instructions. As a probe, a 15-nucleotide random DNA oligo, 5’-/56-FAM/TGA AGT AAT CTG TTA /3BHQ_1/-3’ labeled with a 5′ FAM and a 3′ black hole quencher (3BHQ) was added to the in vitro transcription-translation (IVTT) mix at a final concentration of 500 nM. To increase the likelihood of gene expression of genomic DNA fragments, the *E. coli* RNA Polymerase holoenzyme (NEB, #M0551, MA, USA) was supplemented at 5,000–10,000 units per 200 µl emulsion, in addition to the T7 RNA polymerase, present in the kit.

#### Fluorescence-assisted droplet sorting (FADS)

The droplets containing the IVTT reaction mix were sorted by fluorescence-activated droplet sorting (FADS) using an On-chip Sort instrument (On-chip Biotechnologies, Japan) in 150m size channel microfluidic chips (Chip-Z1000-w150). As stealth fluid, HFE7500 (3M™ Novec™ 7500 Engineered Fluid) supplemented with 0.1% 008-fluorosurfactant from RAN biotechnologies was used.

The droplets were gated according to FSC/SSC to filter out emulsion artifacts and sorted according to either GFP or Fluorescein (results section 5 and 6, respectively, FL-2, ex. 488 nm, em. 543 ± 22 nm). Laser intensities were empirically adjusted for each case

Given the expected low hit rate, sorting gates were set so that (it’s “so that”) sorted droplets showed a minimum of 3 and 10–fold-change from the main population for GFP and Fluorescein which corresponded to 0.01%; and 0.05% and 0.1% for GFP and Fluorescein selection round 1 and 2, respectively.

#### Microdroplet imaging

To image DNF producing droplets (see above), HFE7500 (3M™ Novec™ 7500 Engineered Fluid) supplemented with 0.1% 008-fluorosurfactant from RAN biotechnologies and 10X SYBR gold was prewarmed at 37°C to help SYBR solubility and 5-10 µl of DNF-generating polydisperse emulsion was added to the mix. DNF-generating droplets and IVTT droplets expressing GFP and mCherry (Figure 3B and S5) were imaged using a Zeiss LSM 510 META microscope. IVTT droplets expressing GFP or containing DNF derived from the E. coli genomic library (Figures S8A and S11, respectively) were imaged using a Zeiss ApoTome Axiovert 200M microscope equipped with AxioCam MR camera.

#### Generation of test metagenomic libraries

Test metagenomic libraries to evaluate HRR viability (results section 4) were generated by TA cloning of fragmented eDNA into pMC174 after linearization with XcmI (NEB #R0533).

Test library 1 (Figure 4 and 5) was generated by RCA of existing Dixie hot spring water fosmid-based pooled metagenomic library from the NEB collection (described in Chuzel et al., 2018). RCA was performed using specific oligonucleotides for a minimum of 10h at 42°C. 1 µg of the resulting product was fragmented using a G-tube (Covaris, MA, USA) to generate ∼6 kb DNA fragments that were further end-repaired and A-tailed using the NEBNext^®^ Ultra™ II End Repair/dA-Tailing Module (#E7546). The resulting product was ligated to pMC174 using T4 ligase (NEB #M0202) for a minimum of 10h at 16° C.

**Figure 4.**
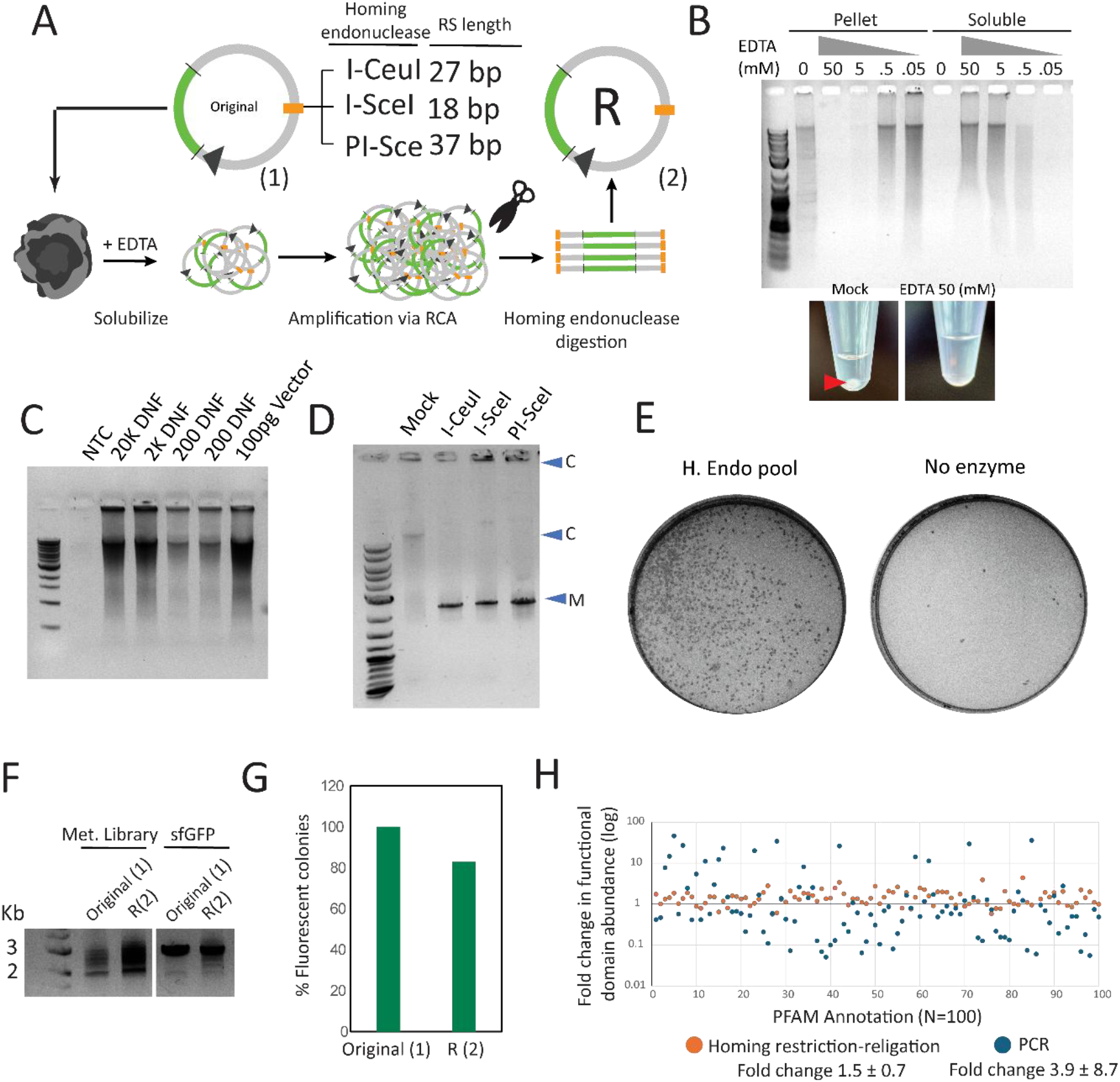
(**A**) Homing restriction–religation (HRR) workflow for reconstituting plasmids from RCA concatemers maintains the functionality of the original vectors. (**B, upper**) Gel electrophoresis of DNFs before and after addition of increasing concentrations of EDTA. Both the soluble fraction and the pellet (after centrifugation) are shown. EDTA treatment shifts DNF material from the pellet into the soluble fraction, indicating solubilization. (**B, lower**) Photographs of the tube bottoms after centrifugation of DNFs, showing that EDTA removes the visible pellet. (**C**) RCA performed with decreasing amounts of solubilized DNF as template. Robust amplification is observed down to ∼200 DNFs, indicating solid amplification sensitivity. (**D**) Restriction analysis of the vector used to develop HRR (pMC174). Digestion with I-CeuI, I-SceI, or PI-SceI converts the original concatemer (C) into linear plasmid species (M) and ligated. (**E**) Plates showing colonies from transformation of 10β bacterial cells with pMC174 either digested with the homing endonuclease pool or left undigested. Colony formation occurs almost exclusively after digestion with the enzyme pool, consistent with reconstitution of viable, monomeric plasmids following cleavage and ligation. (**F**) Agarose gel showing original and reconstituted (R) plasmids from a metagenomic library and a GFP-expressing vector; both pairs run at comparable sizes and display similar migration patterns. (**G**) Functional copy number measured after transformation of either the original or reconstituted GFP vector into an expression strain of E. coli; reconstituted plasmids show a 10–20% decrease in functional copies. (**H**) PFAM domain distribution in the metagenomic library before and after plasmid reconstitution by PCR or HRR; HRR-reconstituted plasmids retain domain frequencies comparable to the original library, whereas PCR introduces severe shifts. For the analysis, the 100 most abundant PFAM annotations were considered.

**Figure 5.**
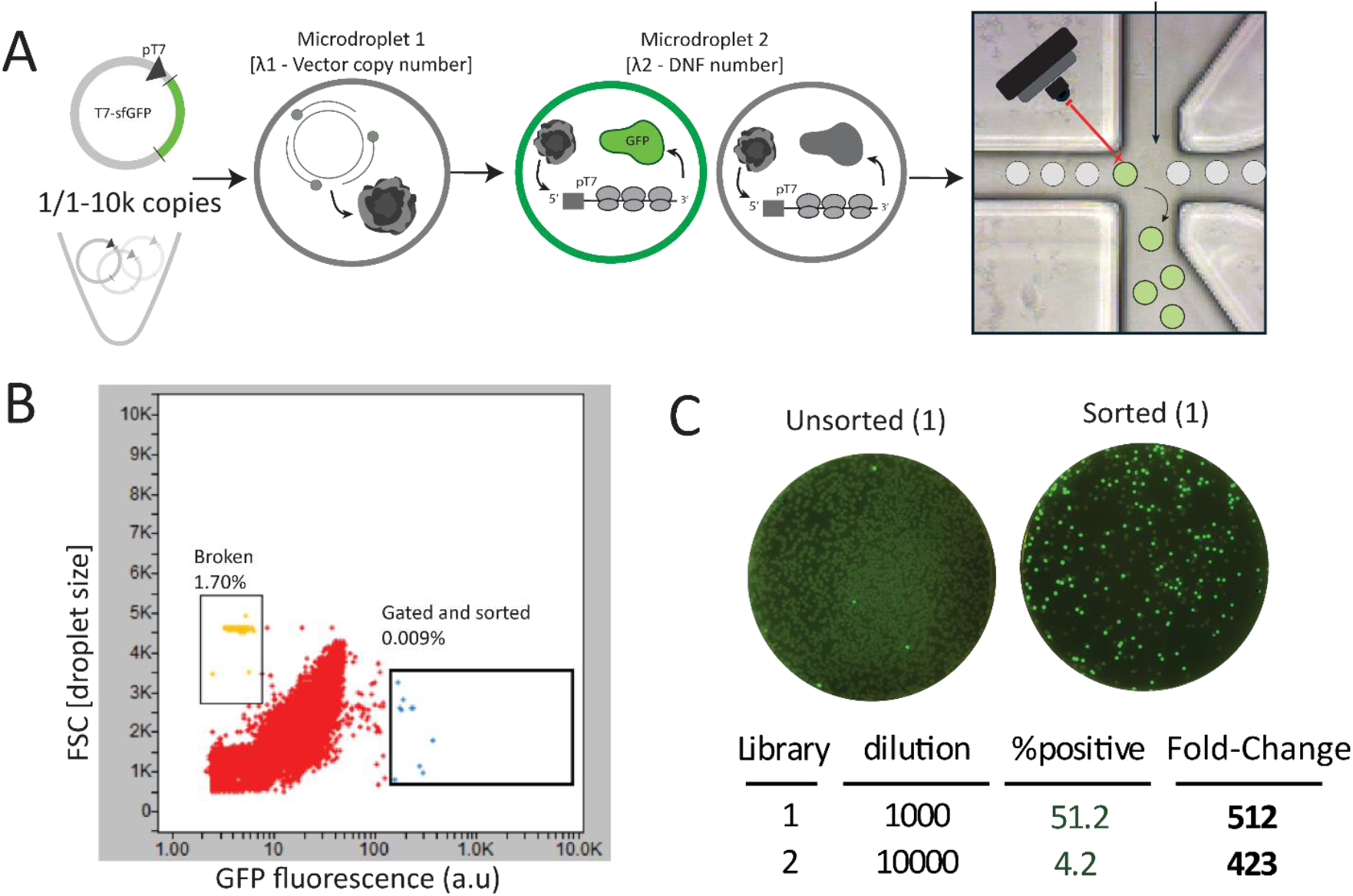
(**A**) Two-step workflow design to assess Ultrahigh Throughout functional selection. A GFP-expressing vector is spiked into a complex metagenomic at high dilution factors and selected by fluorescence by FADS. (**B**) Representative FADS sorting step showing a sorting gate of IVTT droplets generated from library 1 set to select fluorescent droplets ∼10-fold over the main population (0.009%). Forward scatter (FSC) accounts for microdroplet size. (**C**) Positive-selected are transformed into an expression strain after HRR. The percentage of fluorescent colonies is indicative of the efficiency of the protocol’s functional enrichment.

Test library used in Figure 4D, was generated as described in Florez-Cardona et al., 2025. In brief, eDNA was extracted from dog stool using the *Quick*-DNA Fecal/Soil Microbe Microprep Kit (#D6012, Zymo Research, CA, USA). 1 µg of extracted DNA was fragmented using a G-tube (Covaris, MA, USA) to generate ∼6 kb DNA fragments. Fragmented DNA was blunted using the Quick Blunting™ Kit from NEB (#E1201) and blunt-ligated into pMiniT2.0 using NEB’s PCR cloning kit (#E1202) following the manufacturer’s instructions.

For both test libraries, the ligation products were purified using the Monarch^®^ PCR & DNA Cleanup Kit (T1130) and electroporated into NEB^®^ 10-beta Electrocompetent *E. coli* (C3020). Resulting colonies were scraped from agar plates after overnight growth and selection.

#### Generation of *E. coli* genomic library

To construct the *E. coli* expression library for functional genomics, gDNA was extracted from 10-beta High Efficiency Competent *E. coli* (C3019) using the NEB’s Monarch^®^ Spin gDNA Extraction Kit (T3010). 1 µg of extracted DNA was fragmented using a G-tube (Covaris) to generate ∼6 kb DNA fragments. To filter out DNA fragments too small to contain a functional ORF, fragmented DNA was purified using NEBNext Sample Purification Beads (#E7104) at 0.65X. The resulting DNA fragments were further end-repaired and A-tailed using the NEBNext^®^ Ultra™ II End Repair/dA-Tailing Module (#E7546). The resulting product was ligated to pMC174 using T4 ligase (NEB #M0202) for a minimum of 10h at 16 °C. The ligation products were purified using the Monarch^®^ PCR & DNA Cleanup Kit (#T1130), electroporated into NEB^®^ 10-beta Electrocompetent *E. coli* (C3020) and selected in LB medium containing Kanamycin for 6h at 37° with constant agitation.

#### Nanopore sequencing and data analysis

Plasmid libraries were sequenced using Oxford Nanopore technology. Libraries were prepared with the transposase-based ONT Rapid Barcoding Kit 96 V14 (#SQK-RBK114.96) using Kit 14 chemistry, pooled, and loaded onto R10.4.1 MinION/GridION flow cells (#FLO-MIN114) on a GridION Mk1 sequencer. Runs were carried out for 24 h, base-calling was performed with Dorado, and post-sequencing analysis was conducted using the EPI2ME platform.

Nanopore reads were annotated with MetaGeneMark for gene prediction, HMMER3 for domain annotation, and the Pfam database for functional classification. Functional domain abundance was calculated as the percentage of each Pfam annotation count relative to the total number of contigs per library. Functional enrichment was assessed as the fold change in domain abundance before versus after each selection round. Domains showing ≥20-fold enrichment relative to the original library were considered significant. Because only ∼10^2^ droplets were recovered per selection round and RCA in the HRR workflow can produce nonspecific amplification (Figure 4A), a minimum cutoff of 0.005% PFAM annotations per read was applied. Full results are provided in Supplementary Data S2.

## Results

### 1 Microfluidics-free generation of DNA nanoflowers (DNF)

To generate clonal DNA templates for efficient protein expression in microdroplets, we first attempted amplifying DNA via RCA and synthesizing proteins in a one-pot reaction. We observed clear inhibition of RCA with increasing concentrations of IVTT mix, in particular upon the addition of the IVTT fraction containing tRNAs, amino acids, and small molecules (Figure S1). A strong inhibition of protein synthesis was observed when RCA components were added to the IVTT mix, with the nucleotide mix and RCA buffer being the strongest inhibitors. This is consistent with previous reports (Seo & Ichihashi, 2023).

In order to circumvent the limitations of DNA amplification and protein synthesis in a one-pot reaction, we developed a microfluidics-free workflow that produces DNA nanoflowers (DNF) from single DNA copies as the first step of a two-step workflow. DNA nanoflowers are self-assembled DNA microstructures formed by prolonged rolling circle amplification (RCA) of a circular DNA template, which yields long tandem-repeat single-stranded DNA that co-condenses with inorganic precipitates into dense, multilayered, flower-like architectures (Galinis et al., 2016; Kim et al., 2018; Li et al., 2023). During RCA, each dNTP incorporation releases pyrophosphate (PPi), which complexes with divalent cations such as Mg^2+^ to form insoluble metal–pyrophosphate (Mg_2_P_2_O_7_) nanocrystals that co-assemble with the repetitive ssDNA and provide the inorganic scaffold for nanoflower formation (Figure 1A,B; Galinis et al., 2016).

With random hexamer primers in RCA reactions, DNFs containing double-stranded DNA concatemers can be produced. These DNFs are fully compatible with in vitro transcription and translation (IVTT) (Figure 1C). Furthermore, when comparing IVTT kinetics between GFP/mCherry reporter plasmids and DNFs derived from the same plasmids, expression from 6.50 × 10^3^ purified DNFs was comparable to, or exceeded, that from 5.60 × 10^3^ plasmid copies (Figure 1C). Therefore, consistent with previous reports (Galinis et al., 2016; Zubaite et al., 2017), our purified DNA nanoflowers retained at least ∼10^4^ functional GFP/mCherry copies that can be efficiently used as templates for IVTT (Figure 1C).

To generate clonal DNA nanoflowers (DNF) from single genes, we developed a simple bulk emulsification protocol that enables parallel processing of plasmid libraries without microfluidic devices. Polydisperse emulsions are produced by vortexing the RCA reaction mix containing a plasmid library diluted so that each droplet contains the desired plasmid copy number according to the Poisson distribution (λ1). Droplet size is tuned by adding a single 3 mm glass bead to a 1.5 mL tube to increase collision frequency, yielding microdroplets with a maximum diameter of ∼40-50 µm (Figure S3). Using this approach, most droplets contain a single DNF after RCA incubation (Figure 2A, B), which facilitates downstream processing and reduces DNF redundancy. Emulsions are then destabilized, DNFs are recovered by centrifugation and extensively washed to remove loose soluble DNA precipitates (Figure 2A), producing a particle suspension in which the vast majority of DNA is stably incorporated into DNFs (Figure 2C, S4).

To confirm clonality, we used a two-step workflow. In the first step, an emulsion produced DNFs at λ1. These DNFs served as DNA templates in a second emulsion containing IVTT reagents at λ2 (Figure 2A). As a reporter, we prepared a library with equimolar GFP and mCherry plasmids. The initial plasmid dilution was set to yield λ1 = 0.5, and DNFs were generated as described in Figure 1A. Purified DNFs were re-encapsulated into 50 µm monodisperse droplets containing IVTT reagents (Figure 2B). The number of DNFs per droplet was adjusted to achieve λ2 = 0.2 or λ2 = 1. After incubation, this produced four distinct droplet populations: green (GFP only, clonal), red (mCherry only, clonal), yellow (both GFP and mCherry, polyclonal), and empty (no protein expression). Each population was enumerated and compared to the expected frequencies from Poisson distribution for the corresponding λ. Observed and expected frequencies matched closely. As a control, λ1 was increased to 3. At λ1 = 3, DNFs were expected to be polyclonal and contain both GFP and mCherry. DNFs produced at λ1=3 yielded yellow droplets regardless of λ2 (Figure S5). These results indicate that our protocol reliably produces clonal DNFs (Figure 3)

### 2 Reverse DNF back to plasmid monomer using Homing restriction-religation (HRR)

One advantage of ultrahigh-throughput screening is the ability to perform sequential, iterative functional selection. DNA fragments that encode the desired catalytic activity can be enriched over multiple rounds, with the output of one round serving as the input for the next. Successful protein-evolution workflows typically recover only small amounts of nucleic acid from selected microdroplets; therefore, a PCR amplification step is usually required to obtain sufficient material for sequencing and for recloning selected inserts into expression vectors for subsequent rounds. When libraries contain inserts of heterogeneous length and base composition, PCR introduces strong biases. Shorter fragments and templates with particular GC content (Benjamini & Speed, 2012) are preferentially amplified (Figure S6A), causing progressive loss of diversity and yielding a library that no longer represents the true selected population (Figure S6A).

Aiming to integrate DNA nanoflowers (DNF) and *in vitro* protein expression into genomic and metagenomic screening pipelines, we developed Homing Restriction-Religation (HRR) to reconstitute functional plasmids from RCA concatemers without introducing amplification bias. We used homing endonucleases, site-specific double-strand DNA endonucleases that cleave long, rare recognition sites. Recognition sites for I-CeuI, I-SceI, and PI-SceI, with lengths of 27, 18, and 37 nucleotides respectively, were introduced into the backbone of our expression vector (Figure 4A). Combining all three sites minimizes the probability that these sequences occur randomly within our libraries.

After microdroplet recovery and destabilization, selected DNFs were solubilized with EDTA, producing a concentration-dependent shift in DNA content from the insoluble pellet to the soluble fraction (Figure 4B). The solubilized DNA was subsequently reamplified by RCA to yield sufficient material for efficient digestion. We successfully reamplified material from as few as 200 DNFs (Figure 4C). The RCA-generated concatemers were then digested with homing endonucleases (Figure 4D), generating monomeric fragments that self-ligated to regenerate functional plasmid monomers (R vector; Figure 4E).

This protocol was tested on two systems: a GFP-expressing vector and a control metagenomic library cloned into the same vector backbone (see Methods). In both cases, the reconstituted R vectors were indistinguishable from the originals by agarose gel electrophoresis (Figure 4F). The ability of HRR to reconstitute functional plasmid monomers was evaluated by transforming both original and R-plasmids encoding GFP into an E. coli expression strain. After HRR, we observed a modest 10–20% reduction in the number of fluorescent colonies (Figure 4G, Figure S7). Analysis of PFAM domain frequencies in the metagenomic library after nanopore sequencing and functional annotation revealed that HRR preserved the original domain distribution (1.5 ± 0.7 average fold-change), whereas PCR reconstitution introduced substantial deviations (1.5 ± 0.7 vs. 3.9 ± 8.7 average fold-change for HRR and PCR, respectively) (Figure 5H, Supplementary Data 1). Together, these results indicate that HRR efficiently reconstitutes plasmid monomers from RCA concatemers with minimal loss of function while largely maintaining the original library diversity.

### 3 Ultrahigh-speed functional selection of compartmentalized *In vitro* reactions in water-in-oil microdroplets

Next, we assessed the pipeline’s capacity for functional selection from complex libraries at ultrahigh throughput by sorting microdroplets based on fluorescence intensity using fluorescence-assisted droplet sorting (FADS). A target GFP plasmid was spiked into a test metagenomic library at 0.01% and 0.001% abundance, generating library 1 and library 2, respectively. These mixed libraries were used to generate DNFs, which were subsequently purified and re-encapsulated into 50 µm IVTT microdroplets (Figure 5A). To adapt the pipeline for functional metagenomics, where hit rates are typically extremely low, λ1 was set to 3 to produce polyclonal DNFs, thereby enhancing sequence diversity in the screen (see 2, Figure S5). λ2 was set to 0.2 for library 1 and 0.5 for library 2. Following incubation for protein expression, IVTT droplets were sorted via FADS using a 150 µm channel chip (Figure 5A). Approximately 930,000 and 750,000 droplets were gated and sorted for library 1 and library 2, respectively, corresponding to 0.007% of each library. Droplets exhibiting GFP fluorescence levels approximately 10-fold higher than the main population were selected at peak sorting speeds of 500 events per second (Figure 5A, B, S8).

Recovered droplets underwent HRR, reconstituted vectors were transformed into an *E. coli* expression strain, and selection efficiency was evaluated by counting fluorescent colonies. Library 1 and library 2 yielded 51.4% and 4.2% fluorescent colonies, corresponding to 512-fold and 423-fold functional enrichment, respectively (Figure 5C, S7).

### 4 Functional genomic screening of E. coli ssDNA interactors

As a demonstration of our pipeline’s capacity to perform unbiased screens for specific enzymatic activities, we designed a proof-of-principal experiment that exploits two advantages of a reconstituted IVTT system: reduced background from having only minimal reaction components and the ability to screen cytotoxic proteins, including DNA-modifying or DNA-interacting enzymes. To identify ssDNA-binding proteins or ssDNA nucleases, we used a 15-nt single-stranded DNA substrate labeled with a 5′ fluorescein and a 3′ quencher (Figure 6A). In the absence of ssDNA binders, the oligonucleotide is flexible, and the quencher suppresses fluorescence. Upon cleavage or protein binding the oligonucleotide adopts an extended conformation that separates fluorophore and quencher, restoring the fluorescein signal. As shown in fig. S8, the fluorescence of the probe increased when ET-SSB (Extreme Thermostable Single-Stranded DNA Binding Protein) is added to the reaction. As a target library, we generated an expression library by fragmenting the *E. coli* genome into 6-10 kilobase inserts that were cloned into our system’s compatible plasmid backbone (Figure 6B, see 2). The library was sufficiently diverse to comprehensively cover the *E. coli* genome (Figure S9).

**Figure 6.**
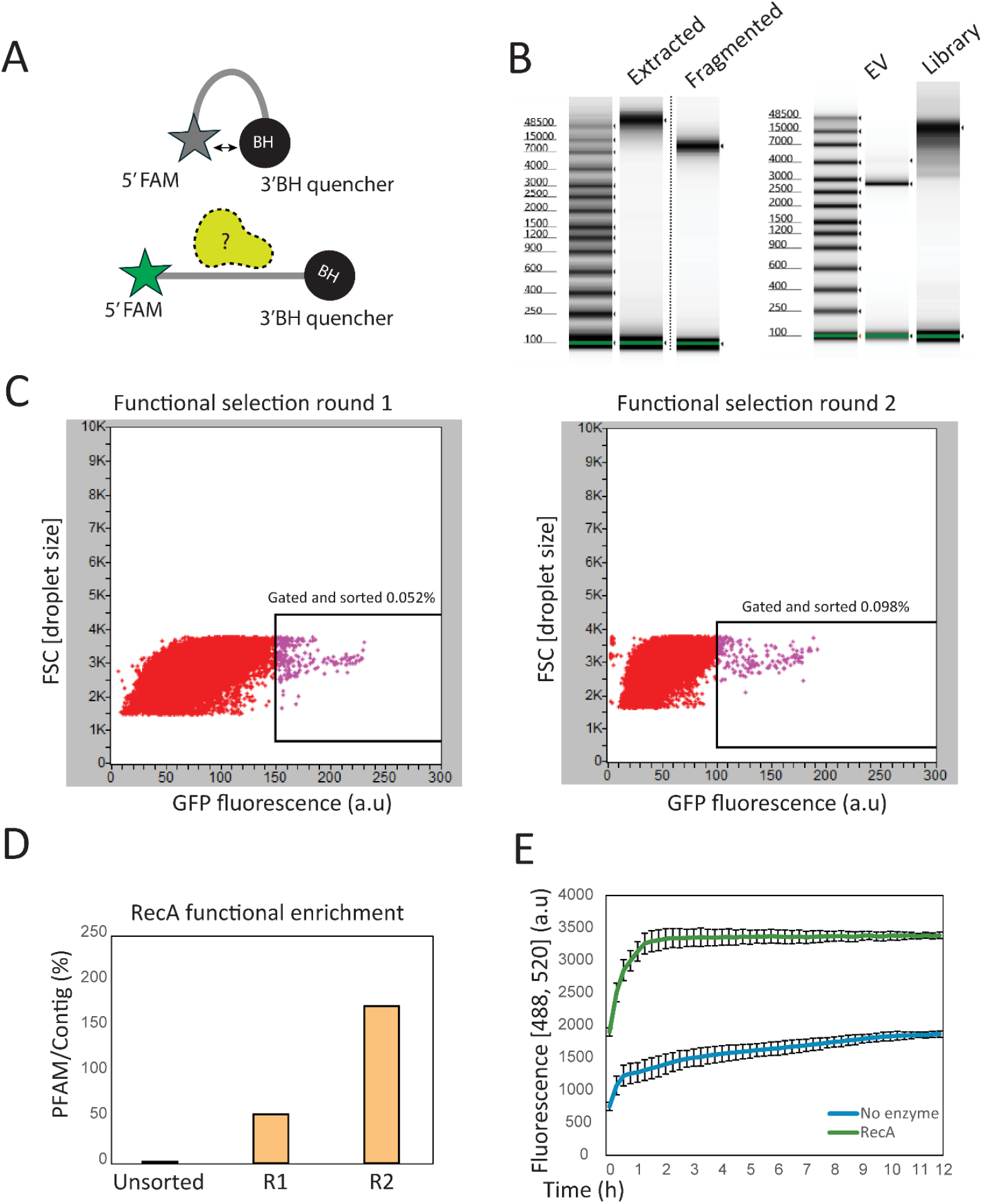
**(A)** A 15-nucleotide single-stranded DNA (ssDNA) substrate labeled with a 5′ fluorescein and a 3′ quencher serves as a reporter for ssDNA-binding proteins. Upon protein binding, the substrate transitions from a quenched, intramolecularly folded conformation to an extended state that spatially separates the fluorophore and quencher, thereby restoring fluorescein fluorescence. **(B)** A genomic library from *E. coli* was constructed by mechanical fragmentation and cloning of 6–10 kb inserts into the pMC174 backbone (EV, empty vector). **(C)** Representative FADS sorting step showing the gating of IVTT droplets generated from DNF derived from the *E. coli* library and supplemented with the 15-nt ssDNA substrate. Microdroplets exhibiting fluorescence levels approximately threefold above the main population were selected, corresponding to ∼0.05% in round 1 and ∼0.1% in round 2. Forward scatter (FSC) reflects microdroplet size. **(D)** Enrichment of RecA Pfam Annotations During Screening. A progressive increase in RecA Pfam annotations per contig was observed, rising from an initial 2% to 229% after two rounds of selection. **(E)** Validation of RecA ssDNA-binding activity. Binding of RecA to the 15-nt oligonucleotide results in a gradual increase in fluorescence.

In plate-based functional screening, genes are heterologously expressed using the endogenous *E. coli* transcription and translation machinery. By contrast, the PURExpress expression system relies on a T7 promoter encoded in the vector backbone. This design restricts the effective sequence space to inserts cloned in the correct orientation, effectively halving the usable library. Even when oriented correctly, random fragmentation places unknown ORFs at variable and often suboptimal distances from the promoter and ribosome binding site.

To address these expression-related limitations and enhance the probability of protein production from individual genes in PURExpress IVTT microdroplets, we adopted a dual-polymerase strategy. In this approach, the T7 RNA polymerase provided in the IVTT kit was supplemented with purified *E. coli* RNA polymerase holoenzyme (core enzyme saturated with the σ70 factor). The functionality of the *E. coli* RNA polymerase holoenzyme in reconstituted cell-free systems has been previously demonstrated (Asahara & Chong, 2010). By enabling transcription from both the T7 promoter and endogenous bacterial promoters present on inserts, this strategy allows gene expression in either orientation. Consequently, a greater fraction of library members can be functionally expressed.

Following the criteria in Section 3, we performed two functional selection rounds on DNFs generated from the *E. coli* genomic library, setting λ1 = 3 to expand the effective sequence space and fixing λ2 = 0.5. In rounds 1 and 2, approximately 1.2 million and 340,000 droplets were gated, respectively. Droplets exhibiting fluorescein signals ≥3-fold above the main population (≈0.09% in round 1 and ≈0.1% in round 2) were sorted at ultrahigh speed and collected as candidate carriers of DNA encoding ssDNA-binding proteins (Figure 6C, Suppl. Data S2). Reconstituted vectors were recovered by HRR after each selection, and resulting plasmids were sequenced using Nanopore technology with functional domain annotation against the Pfam database. Hits were defined using a minimum enrichment threshold of 20-fold. Functional annotation revealed strong enrichment of a contig encoding the recombinase RecA (Figure 6D, Table S1), a protein that binds single-stranded DNA and catalyzes ATP-dependent homologous strand exchange during recombination (Roca & Cox, 1990). That contig showed 29-fold and 92-fold enrichment relative to the original library after the first and second rounds, respectively, and contained neighboring ORFs (mltB, pncC, srlA) not evidently related to ssDNA binding (table S1). To validate second-round outputs, plasmids from 10 independent colonies were sequenced; 6 matched the RecA contig identified in bulk sequencing. The IVTT product of three of these plasmids were independently validated for ssDNA-binding activity in IVTT assays (Figure S11), and ssDNA binding was further confirmed with purified RecA protein (Figure 6E).

## Discussion

In this study, we developed a pipeline for functional genomic screening that integrates reconstituted *in vitro* protein expression (PURExpress system) with Ultrahigh-throughput droplet microfluidics (Figure 7). To ensure sufficient protein yield from clonal DNA templates, we adapted the DNA nanoflower (DNF) strategy of Galinis et al. (2016) and established a two-step, segmented workflow that couples DNF amplification with clonal protein expression. In the first step, a polydisperse emulsion amplifies the plasmid library to generate DNFs; these DNFs are subsequently re-encapsulated into a second emulsion containing IVTT reagents. By eliminating specialized microfluidic operations such as droplet merging or picoinjection, this design removes a major bottleneck between DNA amplification and protein expression while maintaining throughput and clonal fidelity.

**Figure 7.**
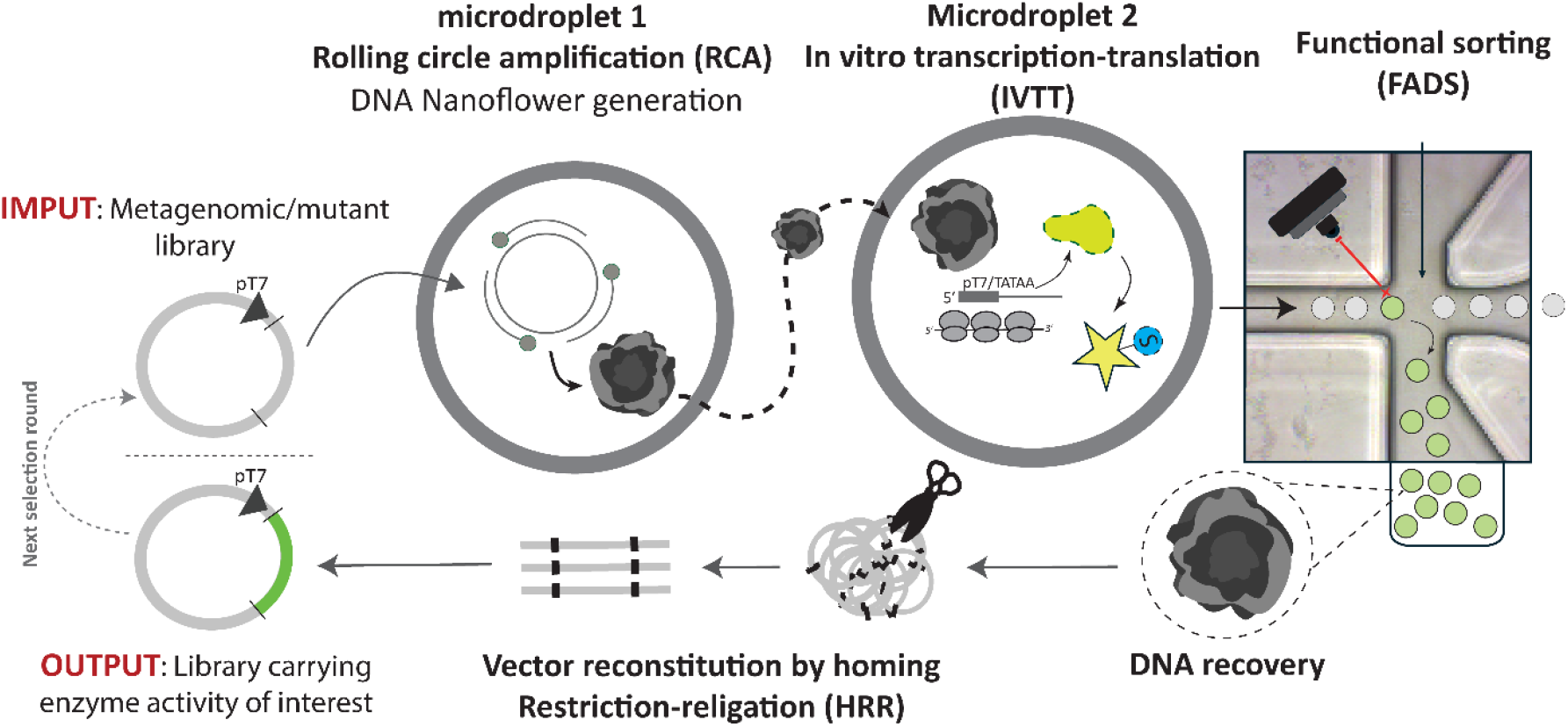
Diagram of the microdroplet-Based Functional Genomic Screening pipeline by combination of DNA Nanoflowers and PurExpress Cell-Free Expression developed in this work.

Our microfluidics-free approach to generating clonal DNFs markedly reduces system complexity, as monodisperse droplets are required only during fluorescence readout to avoid size-dependent signal artifacts. This simplification enables parallel processing of multiple libraries and owing to the long-term stability of DNFs in solution, substantially decreases experimental time while improving reproducibility. Beyond providing clonal gene amplification and enhancing protein expression, the method also permits estimation of DNF concentration by counting SYBR Green-– stained DNFs under a microscope. This capability allows relatively precise control of the Poisson parameter (λ) during subsequent encapsulation steps.

The pipeline also incorporates a module for reconstituting vectors from concatemers generated by rolling circle amplification via homing restriction (HRR), facilitating unbiased and iterative selection rounds. While restriction enzyme-based approaches have shown success in similar contexts (Christ et al., 2006; Fryer et al., 2025; Grasemann et al., 2023), our use of homing endonucleases allows application to unknown DNA sequences. The extremely low probability of encountering their large recognition sites by chance (approximately once per 7 × 10^9^ base pairs according to Jasin, 1996) makes this strategy particularly well-suited for functional genomics and metagenomics. Besided HRR, alternative approaches that could be very interesting to explore include the use of programmable nucleases such as prokaryotic Argonautes or even caspases, as both have demonstrated utility when combined with RCA for nucleic acid detection in molecular diagnostics (Graver et al., 2024; Kropocheva et al., 2022; Zhu et al., 2023). Notably, our HRR results indicate that although RCA is known to produce amplification chimeras (Lasken & Stockwell, 2007), the generation of DNFs does not significantly alter library diversity, at least at the functional level. Non-functional ORFs resulting from amplification artifacts are marginally expected (10–20%) (Figure 4C) but are likely to be eliminated through functional selection rounds. In line with this, our proof-of-principle *E. coli* screening demonstrates that the reconstituted plasmids obtained after multiple rounds of selection are fully functional (Figure S11).

Pilot experiments using GFP as a reporter demonstrated 400–500-fold enrichment when screened by fluorescence-activated droplet sorting (FADS) at ultrahigh speed. A 200 μl emulsion of 50 μm droplets (approximately 2 million microdroplets) can be screened within a few hours. This setup is ideal for assays with broad dynamic ranges (e.g., ∼10-fold signal differences), though it may be less effective for targets with narrower dynamic ranges. Additionally, ultrahigh throughput increases the likelihood of coincident droplet detection, which can elevate the false positive rate, particularly when screening for rare hits.

Our proof-of-principle assay successfully identified and retrieved RecA ssDNA binding activity from the *E. coli* genome. A single round of selection yielded significant enrichment, and a second round produced functional clones ready for validation and expression in downstream assays, without the need for further clone arraying or individual screening. One obvious question raised by these results is why, given the large number of ssDNA-binding and nuclease enzymes encoded in the *E. coli* genome and the acceptable coverage of our library, only a single contig was recovered. Three factors might contribute. First, IVTT expression efficiency is uneven: constructs driven by a T7 promoter and with correctly oriented open reading frames achieve optimal expression, whereas ORFs depending on native *E. coli* regulatory elements are expected to be expressed at lower levels. Second, although IVTT systems are cleaner than cellular lysates, they contain substantial nucleic acids and other components that can compromise target enzyme activity, so not every expressed protein will produce a detectable fluorescent signal. Third, the selection design itself amplifies these biases: multiple rounds and stringent thresholds create a funnel effect that favors high-signal hits and disfavors modest but potentially relevant activities. These limitations are addressable through the pipeline’s intrinsic customizability. We demonstrate that the number of plasmid molecules amplified and embedded into DNFs determines the sequence space that can be screened. This creates a highly customizable system that allows the clonality of DNFs to be tuned to the expected hit rate, reducing the number of DNFs required to cover a library according to its anticipated complexity. Thus, to maximize discovery of broad enzymatic activities, one can use long-insert libraries with moderately polyclonal DNFs to expand the sequence space sampled per round. To capture rare or weak activities, shorter inserts, reduced sorting speeds, and additional selection iterations can improve sensitivity and reduce false positives. Library composition may also be critical, since ORFs placed under T7 control are likely to achieve substantially higher expression than those driven solely by native regulatory sequences. Further successful discovery campaigns are necessary to test these scenarios.

When applied to metagenomics, we anticipate that the dual–RNA polymerase strategy used in this assay will facilitate the expression of bacterial and archaeal genes, depending on their accessibility to the *E. coli* RNA polymerase holoenzyme. Conceptually, this is analogous to traditional functional metagenomic approaches, in which approximately 40% of genes are predicted to be readily expressible in this manner (Gabor et al., 2004).

To the best of our knowledge, this is the first report of successful ultrahigh-throughput functional screening of unknown and complex DNA pools relying exclusively on in vitro protein expression. As demonstrated here, the cell-free approach to functional metagenomics offers a powerful alternative to conventional platforms, addressing several key limitations. First, it enables reliable screening of enzymes that are difficult to assay in complex biological environments such as cell lysates or intact cells, particularly DNA-interacting and DNA-modifying enzymes, which are often inaccessible to traditional workflows. These enzyme classes are of considerable interest and commercial value in the era of synthetic biology. Second, this workflow facilitates the screening of enzymes that are toxic when expressed in *E. coli*, thereby overcoming the in vivo bias that frequently impedes enzyme discovery.

Further development efforts of the method will focus on microdroplet recovery after FADS to minimize microdroplet loss, which has been observed in the course of this current work. Whether HRR can function without additional DNA amplification steps that might facilitate downstream processing steps is yet to be investigated.

Taken together, this study expands the enzyme discovery toolbox by introducing a novel approach that is readily adaptable to customized screening campaigns and diverse laboratory settings. We believe it will enable the identification of new or enhanced catalytic activities in the future.

## Author contributions

J. C. B. data curation; J. C. B formal analysis; J. C. B, A. T., M. S., methodology; J. C. B writing-original draft; J. C. B., M. C., J. B. writing-review and editing; J. C. B, A. T., M. S., O. P. validation; M. C., J. O. supervision; J. C. B., M. C., M. S., O. P, J. B., J. O. conceptualization; M. C., J. O. funding acquisition; J. C. B., A. T., M. C, M. S., O. P., J. B. investigation; M. C., J. O. project administration.

## Acknowledgments

We thank Dr. Donald Comb for research funding at New England Biolabs. We thank the sequencing core of New England Biolabs for their dedicated work. We also thank Léa Chuzel, Sean Lund and Peter Weigele for inspiring the method development with fruitful discussions, and Tom Evans for comments and critical review of the manuscript prior to its submission. Lastly, we thank Muddy, generous donor of eDNA for our test libraries.

