## Supplemental Figures for "Development of a Microdroplet-Based Functional Genomic Screening pipeline by combination of DNA Nanoflowers and PURExpress Cell-Free Expression"

### Suplemental data

SD1: Test metagenomic library – HRR – PCR comparison

SD2: E. Coli screening data

### Supplemental figures

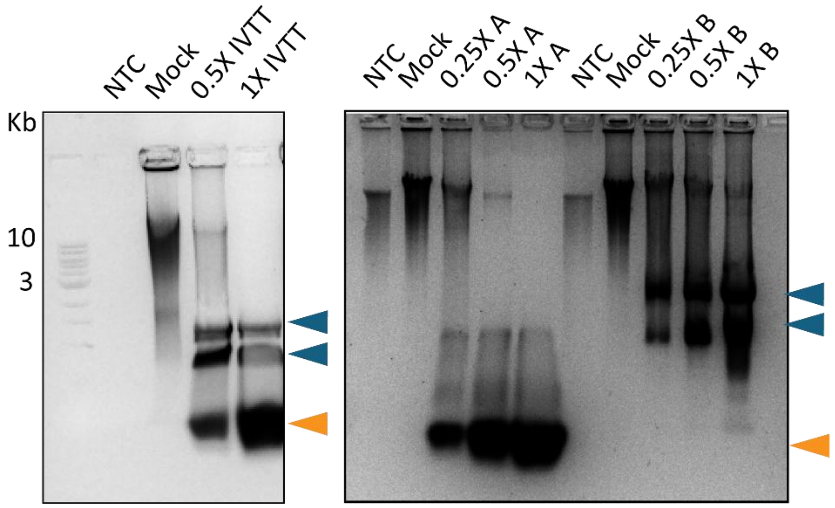


**Figure S1** (**left**). RCA is inhibited by components of the PurExpress IVTT mix. Left: increasing amounts of PurExpress (Solutions A + B) progressively reduce RCA product formation. Lanes: NTC (no template control), Mock (no addition), 0.5X (50% V/V PurExpress), 1X (100% V/V PurExpress). Right: the two PurExpress fractions were tested separately to identify the inhibitory component(s). Solution B (ribosomes, translation enzymes and factors) and Solution A (tRNAs, amino acids and small molecules) were titrated as shown. Lanes: NTC, Mock, 0.25X (25% V/V), 0.5X (50% V/V), 1X (100% V/V). Progressive addition of Solution A causes severe inhibition. Bands corresponding to ribosomal subunits and tRNA in the PurExpress kit are indicated by blue and orange arrows, respectively.

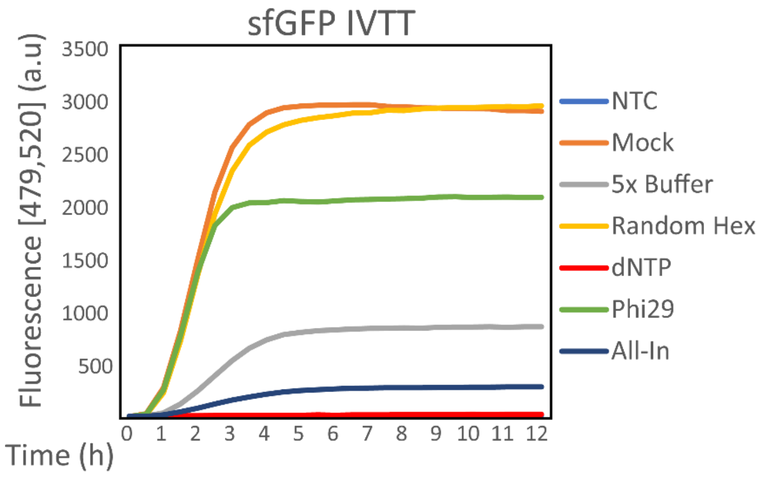


**Figure S2**. Components of the Phi29-XT RCA mix inhibit IVTT. Individual RCA components or the complete mix ("All-in") were added to a PureExpress in vitro translation reaction using 100 ng of PucT7-GFP as template. Addition of Phi29 polymerase and of dNTPs produced the largest inhibition of GFP expression.

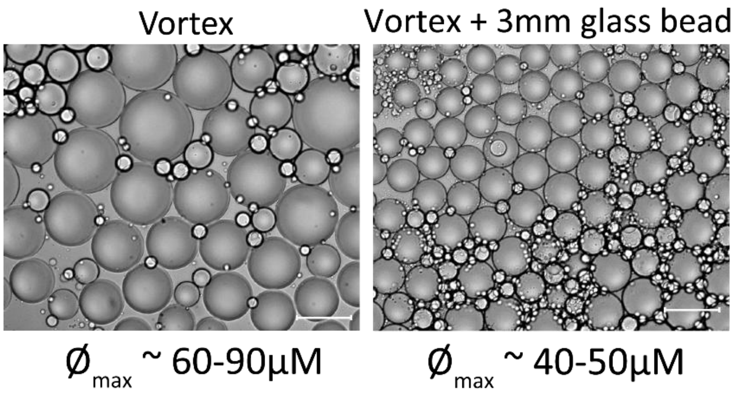


**Figure S3**. Polydisperse water-in-HFE7500 microdroplets generated by vortexing in a 1.5 mL tube. Without agitation aids, droplet diameters were approximately 60–90 µm. Inclusion of a 3 mm glass bead during vortexing reduced the maximum diameter to ~45–50 µm and narrowed the size distribution. Scale bar: 50 µm.

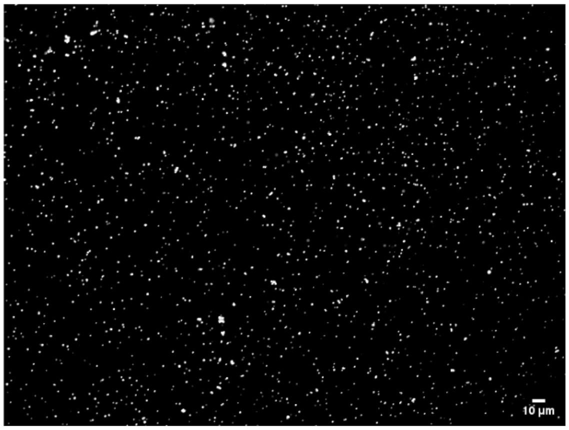


**Figure S4**. DNFs were generated, purified, and stained with SYBR Gold as described in the Methods. The image is a representative micrograph used for particle counting in ImageJ. Scale bar: 10 µm

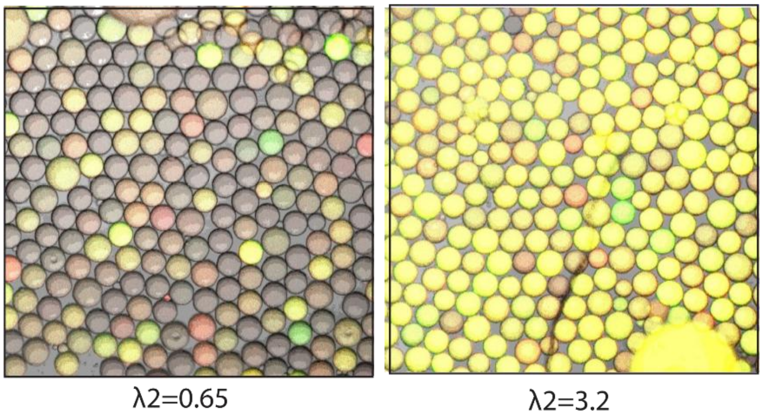


**Figure S5**. Confocal images of IVTT reactions in 50 µm microdroplets used to assess DNF clonality in a two-step workflow (Figure 3). DNFs were generated at λ1 = 3, purified, then re-encapsulated into a second emulsion containing IVTT reagents at λ2 = 0.65 (left) or λ2 = 3.2 (right). Images show representative fluorescence from IVTT expression within individual droplets, used to evaluate clonality of encapsulated DNFs. Increasing λ1 to 3 produces predominantly polyclonal IVTT droplets that coexpress GFP and mCherry (appearing yellow/orange) regardless of λ2.

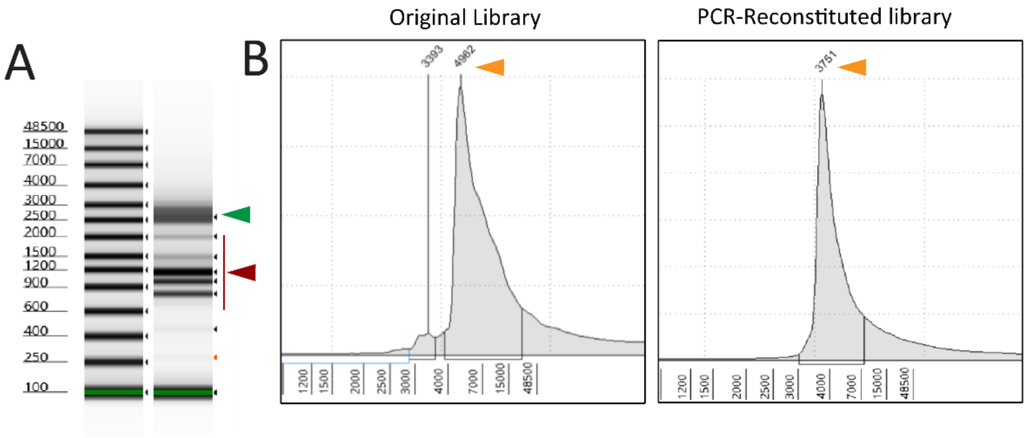


**Figure S6**. (**A**) PCR amplification of inserts from a plasmid metagenomic library exhibits size-dependent bias. TapeStation analysis of the PCR product is shown; the expected average insert size (2.5–3.0 kb) is marked with green arrows, while the predominantly amplified smaller species are marked with red arrows, indicating preferential amplification of shorter inserts. (**B**) TapeStation profiles comparing the original plasmid metagenomic library and the library after PCR amplification of inserts followed by recloning. Size analysis reveals an average reduction of ~1 kb in insert length after the PCR/recloning workflow (main peak indicated with orange arrow).

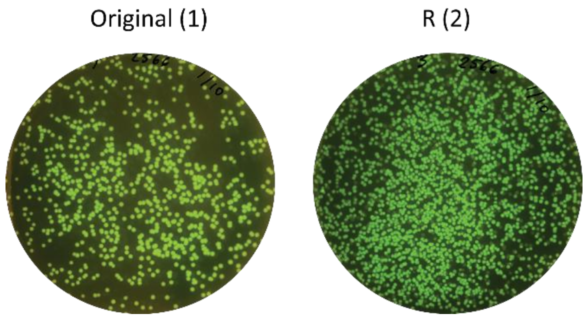


**Figure S7**. Functional copy number measured after transformation of either the original or reconstituted GFP vector into an expression strain of E. coli; reconstituted plasmids show a 10–20% decrease in functional copies. Plates are representative of colony count in figure 5B.

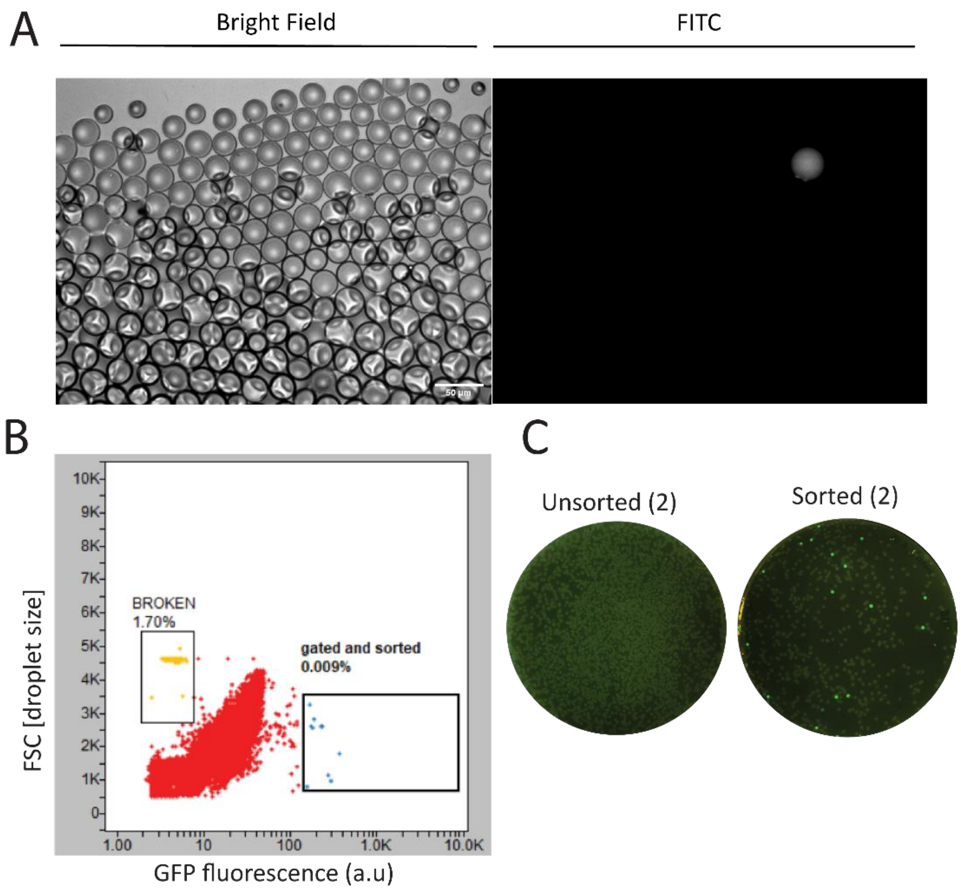


**Figure S8.** (**A**) Bright-field and fluorescence images of 50 µm microdroplets containing DNFs prepared from metagenomic Library 1 (Figure 5), spiked with 0.01% GFP-expressing vector. A single fluorescent-positive droplet is visible, indicating one GFP-producing construct in the droplet population. Scale bar: 50 µm. (**B**) Representative FADS sorting step for IVTT droplets generated from Library 2. A sorting gate was set to select droplets with fluorescence ~10-fold above the main population (target frequency ~0.009%). Forward scatter (FSC) correlates with droplet size. Fluorescent-selected droplets were processed by HRR and transformed into an E. coli expression strain. (**C**) The fraction of fluorescent colonies recovered (4.2%) indicates the protocol's functional enrichment efficiency (≈423-fold enrichment; see Figure 5C).

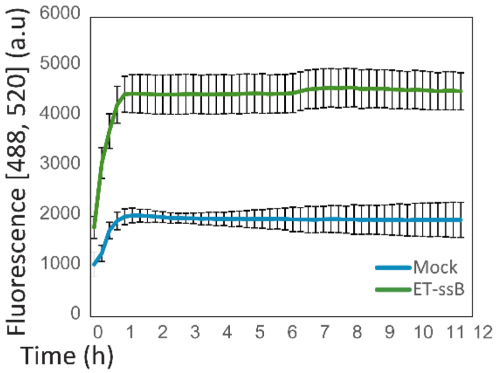


**Figure S9.** Binding of ET-ssB to a 15-nt oligonucleotide labeled with a 5′ fluorescein and a 3′ Black Hole Quencher (see Figure 6A) produces a progressive increase in fluorescence as binding separates the fluorophore from the quencher.

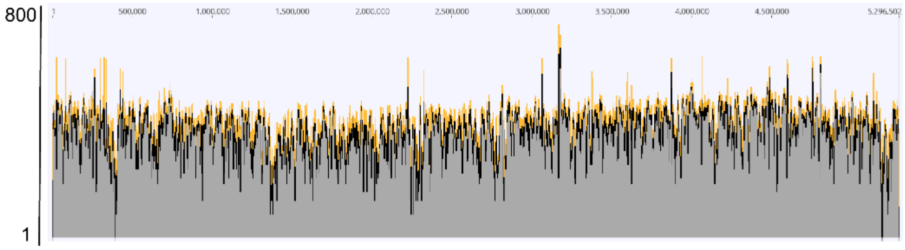


**Figure S10**. Coverage of the E. coli genomic plasmid expression library used in this study. The library was sequenced by Nanopore whole-plasmid sequencing and ~100,000 reads were mapped to the E. coli reference genome (U00096) using minimap2, generating the coverage profile shown.

| PFAM domain | Gene | description | Abundance (PFAM/contig, %) | Fold-Change |
|---|---|---|---|---|
| RecA | RecA | recA bacterial DNA recombination protein | 229.76 | 92.88 |
| SLT_2 | mltB | Transglycosylase SLT domain | 190.19 | 150.92 |
| CinA | pncC | Competence-damaged protein | 120.70 | 109.11 |
| EII-GUT | srlA | PTS system enzyme II sorbitol-specific factor | 94.44 | 145.32 |
| RecA_C | RecA | RecA C-terminal domain | 65.79 | 79.74 |

**Table S1**. Pfam annotation of the contig enriched for ssDNA-binding activity after two functional selection rounds. Abundance is reported as the ratio of Pfam annotations to the total number of contigs sequenced by Nanopore (in %). Foldchange is calculated relative to the input E. coli genomic library.

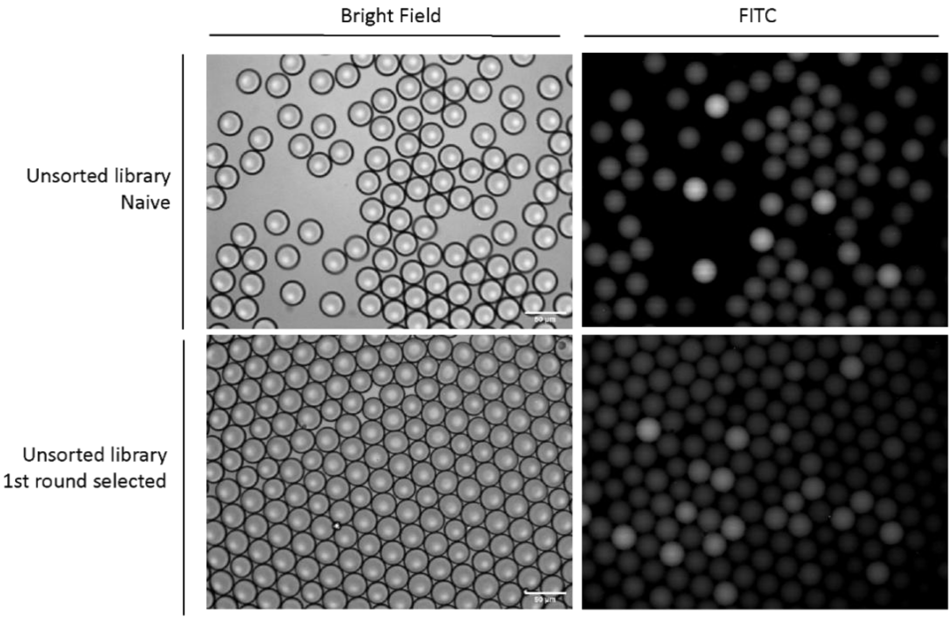


**Figure S11.** Bright-field and FITC images of IVTT monodisperse microdroplets containing DNFs derived from the first and second rounds of functional selection of the E. coli library for ssDNA binders. DNFs were produced with $\lambda 1 = 3$ in both rounds and re-encapsulated for IVTT at $\lambda 2 = 0.5$. Images show representative fields from each round; an increase in the number of FITC-positive droplets is observed in the second round compared with the first, consistent with enrichment of ssDNA-binding activity.

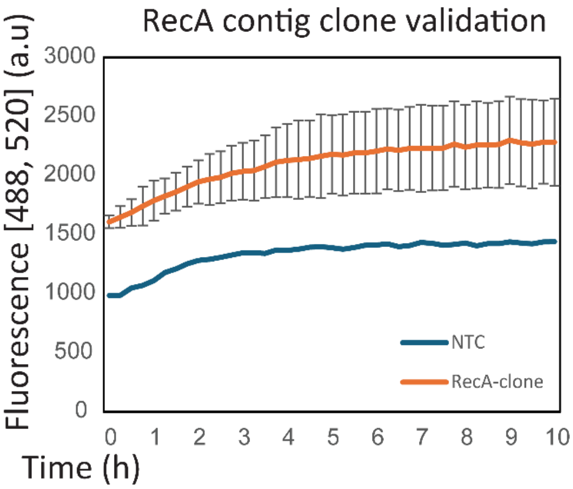


**Figure S12.** Validation of three RecA-positive colonies isolated from the second round of the E. coli screen. IVTT reactions were performed in bulk under conditions matching the screening workflow using each colony as input. Resulting products were assayed for ssDNA-binding activity in buffer containing 70 mM Tris-HCl, 10 mM MgCl2, and 5 mM DTT. Results confirm activity of the RecA-positive clones under the assay conditions described.
